# Remdesivir, Zidovudine (AZT) and Nevirapine inhibit Chandipura virus replication through high energy interactions with the RdRp domain of the polymerase protein L

**DOI:** 10.1101/2022.03.02.482698

**Authors:** Dibyakanti Mandal, Deeksha Pandey, Debi P. Sarkar, Manish Kumar

## Abstract

Chandipura Virus (CHPV), a rhabdovirus belonging to mononegavirales, is an emerging pathogen in Indian subcontinent. The virus infection causes fever, brain encephalitis among the young children below 14 yrs of age. In recent past, several outbreaks and deaths among children were reported from in India. There are no targeted drugs or vaccines available against CHPV and symptomatic treatments are the only option. In this background, we aimed to investigate the inhibitory effects of some priviously known viral RNA polymerase inhibitor drugs on CHPV replication. First, we examined remdesivir, which is known to inhibit HCV, Ebola and SARS-CoV-2 replication and close structural similarity along with conserved residues in the finger region of RNA dependent RNA polymerase (RdRp) domain is the basis of replication inhibition. Our results showed that remdesivir inhibits CHPV replication in vero E6 cells to a significant level. In this study we have also included non-nucleoside anti-retroviral inhibitor (NNRTI) drug nevirapine, and nucleoside inhibitor (NRTI) drug AZT (Zidovudine) to determine if these are also able to inhibit CHPV replication. Interestingly, we observed inhibition of CHPV replication by both nevirapine and AZT (in the order nevirapine>AZT), albeit to a lesser extent compared to remdesivir. We next performed molecular docking and modeling study to get an insight about the interactions of these drugs with CHPV polymerase protein. Modeling study predicts that remdesivir has most favourable CHPV polymerase binding energy among these three drugs. Both remdesivor and AZT binds near the polymerase active site through interctions with residues in finger and palm regions of RdRp. In contrast, nevirapine binds to the N-terminal domain (NTD) of the RdRp. In summary, we found remdesivir as a potent inhibitor of CHPV. A combination therapy including remdesivir, nevirapine and AZT may be a better drug cocktail to treat CHPV disease. Our findings warrant further studies of these drugs against CHPV in animal models for clinical use in near future.

## Introduction

Viral polymerases are potential drug target because of it’s key role in virus RNA transcription and genome replication. In past three decades detailed information was gained about the structure-function relationship of different families of polymerases isolated and chanraterized from RNA and DNA viruses **(1-3)**. X-Ray crystallographic studies of polymerases with bound inhibitors helped in understanding 3D structure and key amino acid residues involved in inhibitor-polymerase interactions. Based on this knowledge, clinically important drugs were designed and used in successful treatment of several recently emerged deadly viruses, such as EBOV, MERS, SARS, SARS-CoV-2 **(4-7)**.

CHPV is an emerging pathogen of Indian subcontinent and in some of the African countries **(8-10)**. Since first reported in 1965 **(11)**, several major outbreaks were reported from the states of Maharastha, Gujrat, Bihar in India **(12-15)**. CHPV causes fever, brain encephalitis among young children below 14 years of age and the case-fatality rate is more than 50% among the infected children. It is a vector-borne disease, spread by phlebotomus sandfly **(16)** and there are possibilities of major endemics by CHPV in future if there is no preventive therapeutics are available. Currently there is no targeted treatment for CHPV encephalitis, and clinical management is done by fluid and electrolyte balance, paracetamol, diclofenac.

VSV is a close family member of CHPV and both viruses share high degree of sequence homology. Crystal stucture of VSV polymerase has been demonstrated recenly and its RdRp (RNA dependent RNA polymerase) was found structurally similar to other rhabdoviruses and negative stranded virus polymerases **(3)**. There is no crystal stucture of CHPV polymerase (called Large or L protein) is available yet but using in-silico modeling we have determined 3D structre of CHPV polymerase L protein, which predicts close similarity in structure to VSV L. Recently published study results also suggest that non-segmented negative stranded RNA viruses (including Rabies, VSV, HRSV) have similar structural domians and have some conserved amino acid sequences in this region of polymerase protein **(17)**. We hypothesized that polymerase drug remdesivir, a ATP analog, which is a potent inhibitor EBOV and also is in emergency clinical use for SARS-CoV-2, may bind CHPV RdRp and inhibit virus replication **(18-22)**. We also assumed that retroviral NRTI drug AZT and NNRTI drug nevirapine may inhibit CHPV replication because of basic structural similarity shared by viral RNA polymerases having a right-hand like structure with finger, palm and thumb subdomains present. Both AZT and nevirapine are clinically used as HAART regimen agaisnt HIV/AIDS in adults and newborn children. AZT inhibits HIV-1 replication through interaction with residues in dNTP binding pocket of reverse transcriptase, whereas nevirapine binds to the palm and connection regions near the active site. AZT acts as a terminator of newly synthesized nucleotide chain **(23,24**). On the other hand, nevirapine and other NNRTIs binding to HIV-1 RT causes structural distortion causing inhibition to overall polymerase activity **(25,26)**. To examine if these three drugs have any inhibitory effects on CHPV replication we have conducted *ex vivo* infection studies in cultured vero E6 cells and analysed viral proteins and viral RNA transcripts from drug treated and infected cell. To better understand the mechanistic insights, we have also done molecular docking and in-silico modeling studies of the chandipura virus plymerase protein L bound with these drugs.

## Methodology

### Polymerase inhibitor drugs

AZT and nevirapine was obtained from Prof. Sanjeev Sinha, All India Institute of Medical Sciences, New Delhi. Powdered nevirapine was dissolved in DMSO (Stock 100 mM) and AZT was dissolved in deionized water as per solubility data (10 mM stock). Aqueous solution of Remdisivir (Jubilant Generics, Authorzed by Gilead Sciences, Stock is 5.2mg/ml or 8.5 mM) was used in our study. 1mM and 100 µM final concentrations of each drug were used to determine the inhibitory effects.

### Cells and Virus

Vero E6 cells were used for all the experiments. Cells were maintained in DMEM (Gibco) containing 10% FBS. Chandipura virus Indian strain, stock (Titer ∼1×10^8^/ml) was obtained from Prof. DJ Chattopadhyay lab (University of Calcutta, Kolkata), was used in this study.

### Drug treatment and virus infection

Vero E6 cells were treated with remdesivir, AZT and nevirapine at 1mM and 100 µM concentrations. 2h post treatment cells were washed with PBS twice and then infected with CHPV at 0.005 moi in serum free DMEM media. Cells were washed 2x with PBS and incubated with complete DMEM containing remdesivir, AZT and nevirapie respectively at same concentration mentioned above. 24h post-infcetion cells were harvested. Total proteins extracted from half of the harvested cells, were analyzed by western blot using anti-CHPV N (obtained from Prof. D Chttopadhyay lab) and anti-tubulin nonoclonal antibody (as internal control). Remaining part of the harvested cells were mixed with 0.5 ml Trizol reagent (Invirogen) and harvested at -80 deg until used. Supernatants containing released virus particles were filtered through 0.45 µm filtration unit (Millipore) and harvested at -80 deg, until these were used for Western blot analysis and plaque assay.

### Plaque assay

Cell free supernatants were used for plaque assay to determine the amounts of virus produced in the presence and absence of drugs. For plaque assay 0.75 million vero E6 cells were seeded in 6-well plates and 48h later cells were infected with serially diluted virus-containing supernatants. Virus dilution and infection was carried out in serum free DMEM media. First, viruses added to cells and incubated at 37°C for 2hrs. Cells were then washed with PBS twice and overlayed with 2x DMEM mixed with equal volume of 2% low melting agarose. Plates were then incubated for 24h at 37 deg. To visualize plaque, cells were stained with crystal violet for 2h, then discarded the agarose overlay and finally wells were rinsed with water. T-test was performed do determine the *p* values.

### Western blot analysis

Infeted cell proteins were extracted using cell lysis buffer (150mM NaCl, 1.0% NP-40, 0.5% sodium deoxycholate, 0.1% SDS and 50 mM Tris-cl pH 8.0) containing protease inhibitor cocktail (Sigma). Lysates were mixed with 2x lameilli buffer containing beta-marcaptoethanol and analyzed on 10% SDS PAGE gel. Viral proteins were extracted from cell-free supernatants by mixing with equal volume of 2x lameilli buffer followed by heating at 100°C for 10 min and proteins were separated on 10% SDS-PAGE gel. Transferred proteins from cell lysates or cell-free supernatants on PDVF/nitrocellulose membrane were detected with anti-CHPV N antibody. Cell lysates were also blotted with anti-beta tubulin antibody (as internal control). Levels of viral proteins present in drug treated and untreated cells were compared.

### Analysis of viral transcripts by PCR and quantitative real-time PCR

For analysis of viral RNA, total RNA from infected cells were isolated using Trizol reagent and following the respective protocol. cDNAs were prepared from 0.5 µg RNA samples using high-capacity cDNA synthesis kit (Applied Biosystems) following standard protocol. cDNA were then used as template to amplify CHPV G (envelope) gene using CHPVG-F (Sequence 5’-CCAAGCTTATGACTTCTTCAGTGACA-3’) and CHPV G-R (5’-CCGTCGACGATATCACTCATACTCTGGCTCTCAT-3’) following standard protocol. Condition for PCR amplification was 1 cycle at 94 deg for 5 min, 25 cylces of 94 deg 15 sec, 55 deg 30 sec and 72 deg 2 min, 1 cycle at 72 deg for 5 min. GAPDH forward and reverse primers (5’-ACCTGACCTGCCGTCTAGAA-3’ and 5’ TCCACCACCCTGTTGCTGTA-3’) were used to amplify GAPDH sequence as internal control.

To get a quantitative measurement of the relative amounts of transcripts synthesized in the infected cells in presence or absence of drugs, we have conducted a SYBR Green based real-time PCR assay. cDNA systhesised were used as template for quantitative PCR. GAPDH and CHPV G specific primers mentioned above were used for this purpose. Real-time PCR were conducted in presence of 1x SYBR Green PCR mix (Applied Biosystems), 200 nM each of forward and reverse primers and 2.5 µl cDNA in a total volume of 20 µl. Reaction was done QuantStudio™ 6 Flex Real-Time PCR System (Applied Biosystems). Reaction was conducted at 25 deg 10 min, 40 cycles of 95 deg 30 sec and 60 deg 1 m 30 sec. Ct values obtained were analysed and comparison was made. ΔΔCt and 2^−ΔΔCt^ was calculated and compared **(27)**.

### Molecular docking and in silico modelling

#### A. 3-D structure prediction and validation

As the 3D structure of L protein (2092 aa) of CHPV is not available in the data bank, we used a homology-modeling server in order to generate a 3-D model of the protein of interest. The 3-D structure of CHPV L protein (2092 amino acids) was modeled using the homology-Swiss-model server. The crystal structure of RNA-directed RNA polymerase L-Structure of the Vesicular Stomatitis Virus L (VSV L) protein in complex with phosphoprotein cofactor of Indiana virus strain San Juan protein (PDB ID: 6U1X) was taken as a primary template to model the structure of CHPV L protein. The sequence identity between the two proteins was 59.79%. The best model was selected among all models provided by swiss-model on the basis of global mean quality estimate (GMQE score) and QMEANDisCo global score. The predicted GMQE and QMEANDisCo global score for the final structure was listed in **Table 1**.

**Table 1:** Target-Template details of selected modeled structures.

| Model | GMQE score | QMEANDisCo Global | Amino Acid | Template | Sequence Identity | Description |
| --- | --- | --- | --- | --- | --- | --- |
| Model | 0.70 | $0.75 \pm 0.05$ | 2092 a.a. | 6u1x:A | 59.79% | RNA-directed RNA polymerase<br>LStructure of the Vesicular Stomatitis Virus L Protein in Complex with Its Phosphoprotein Cofactor (3.0 Å resolution) |

The reliability of the modeled structure was assessed using Structural Analysis and Verification Server (SAVES) (http://nihserver.mbi.ucla.edu/SAVES). SAVES calculates the backbone conformation and overall stereochemical quality of the modeled structures by analyzing the phi (Φ) and psi (ψ) torsion angles using PROCHECK **(28)** to cross-check the Ramachandran plot statistics (**Table 2**). The nonbonded atomic interactions were examined using the ERRAT program **(29)**. Prior to docking, the energy of the modeled structure was minimized using steepest descent (100 steps) and conjugate gradient (500 steps) method from SPDB viewer software platform.

**Table 2:** Model validation.

| Model | Ramachandran Plot Statistics |  |  |  |  | ERRAT Statistics |
| --- | --- | --- | --- | --- | --- | --- |
|  | Residues in most favorable region | Residues in additional allowed region | Residues in generously allowed region | Residues in disallowed region | Number of non-glycine and non-proline residues | Score |
| Model | 1658 (89.4%) | 181 (9.8%) | 8 (0.4%) | 7 (0.4%) | 1854 (100%) | 92.75 |

#### B. Protein-ligand docking protocol

The modeled structure of CHPV L protein was docked with three antiviral drugs namely, nevirapine, zidovudine and remdesivir. The molecular coordinates of all three drugs were retrieved in PDB format from the DrugBank database (http://www.drugbank.ca/). The drug bank ID of nevirapine, zidovudine and remdesivir were DB00238, DB00495 and DB14761 respectively. Both protein and ligand structures were pre-processed using AutoDockTools (ADT) version 1.5.6. The ligand preparation was done by adding charges to the hydrogen atom. From the protein molecule polar hydrogen atoms and Kollman charges were added and all non-protein molecules were removed. Finally both pre-processed protein and ligand structures were converted into the PDBQT format using ADT with default parameters. Each drug molecules such as Nevirapine, Zidovudine (AZT) and Remdesivir were docked on modeled CHPV L protein structure using the AutoDock4 with default parameters and the mean predicted binding affinity from the set of predicted binding poses was accepted as the true binding affinity for each docking run. The maximum number of poses per ligand was set to 10. Each docked complex of all the three drugs were also selected on the basis of their lowest binding free energy.

#### C. Validation of docking results

Complexes with lowest binding energy were further used for estimating the consensus scoring function X-score V2.1. X-Score calculates the binding energy (kcal mol−1) of the ligand to the protein as negative logarithm of the dissociation constant of ligand, −log (*K*d). X-score is basically the average of three empirical scoring functions that includes terms for hydrogen bonding, van der waals interaction, deformation penalty and hydrophobic effects. LIGPLOT v4.4 was used to find the interacting residues/atoms between the CHPV L protein and nevirapine, AZT and remdesivir. Also, two output files are written where the hydrogen bonded interactions and non-hydrogen bonded interactions are tabulated. All molecular images of interaction were generated through PyMOL version 2.0.

## Results

### Remdesivir inhibits CHPV replication and virus production in vero E6 cells

To determine the effects remdesivir on CHPV replication, we treated cultered monolayer of vero E6 cells with 1 mM and 100 µM final concentrations of the drugs. Untreated cells were taken as control for comparison. After 2h drug treatment cells were infected with CHPV as described in the methodology. 24h post infection cells were viewed under microscope, cells and supernatants were harvested. Microscopic images (bright field) showed visible cytopathic effects of CHPV infection on drug unterated cells and cells were rounded (Fig. 1A, middle). Drug treated but uninfected cells were healthy, suggesting that remdesivir did not affect cell growth or morphology (Fig 1A, top). Interestingly, we found that cells that were treated with 1 mM remdesivir and infected with CHPV had no or a minimum cytopathic effects, suggesting significant supression of virus replication by remdesivir (Fig. 1A, bottom). To confirm this results, we have analyzed 1 mM and 100 µM remdesivir treated and CHPV infected cell proteins by western blot analysis. Blotting with anti-CHPV N antibody showed that compared to drug untreated cells, levels of N proteins in 1 mM remdesivir treated cells were much lower. Lesser amounts of N levels were also observed in 100 µM drug treated cells (Fig. 1B, top). We further analyzed virion proteins obtained from cell-free supernatants. We observed significant reduction in the levels of virus production in 1 mM remdesivir treated vero cells and moderate decrease in 100 µM remdesivir treated cells (Fig. 1B, middle). We looked at cellular β-tubulin protein as an internal control. Blot with anti-β-tubulin antibody confirmed that low CHPV N protein in drug trated cells are not because of lower total cell proteins used for analysis (Fig. 1B, bottom).

**Figure 1:**
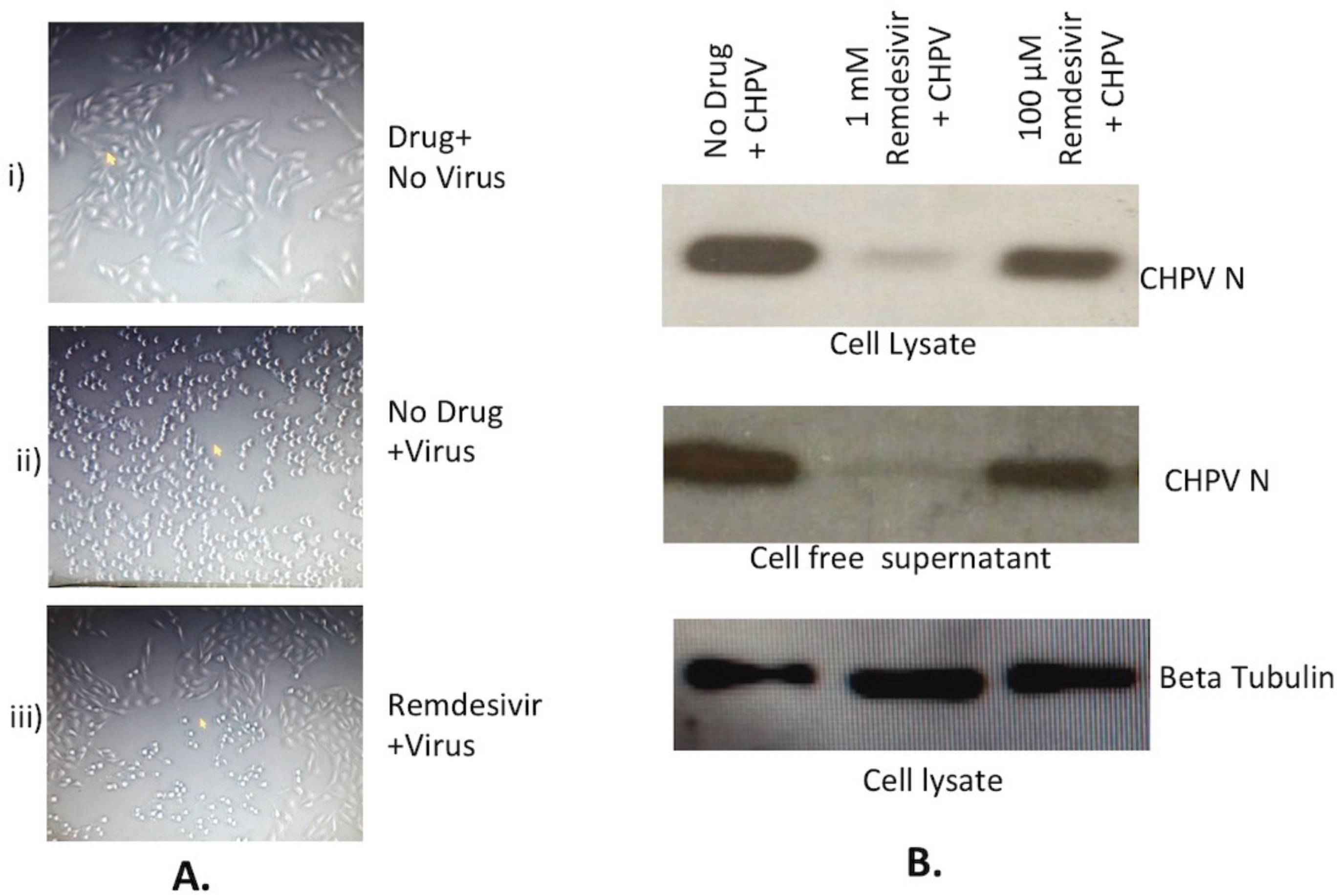
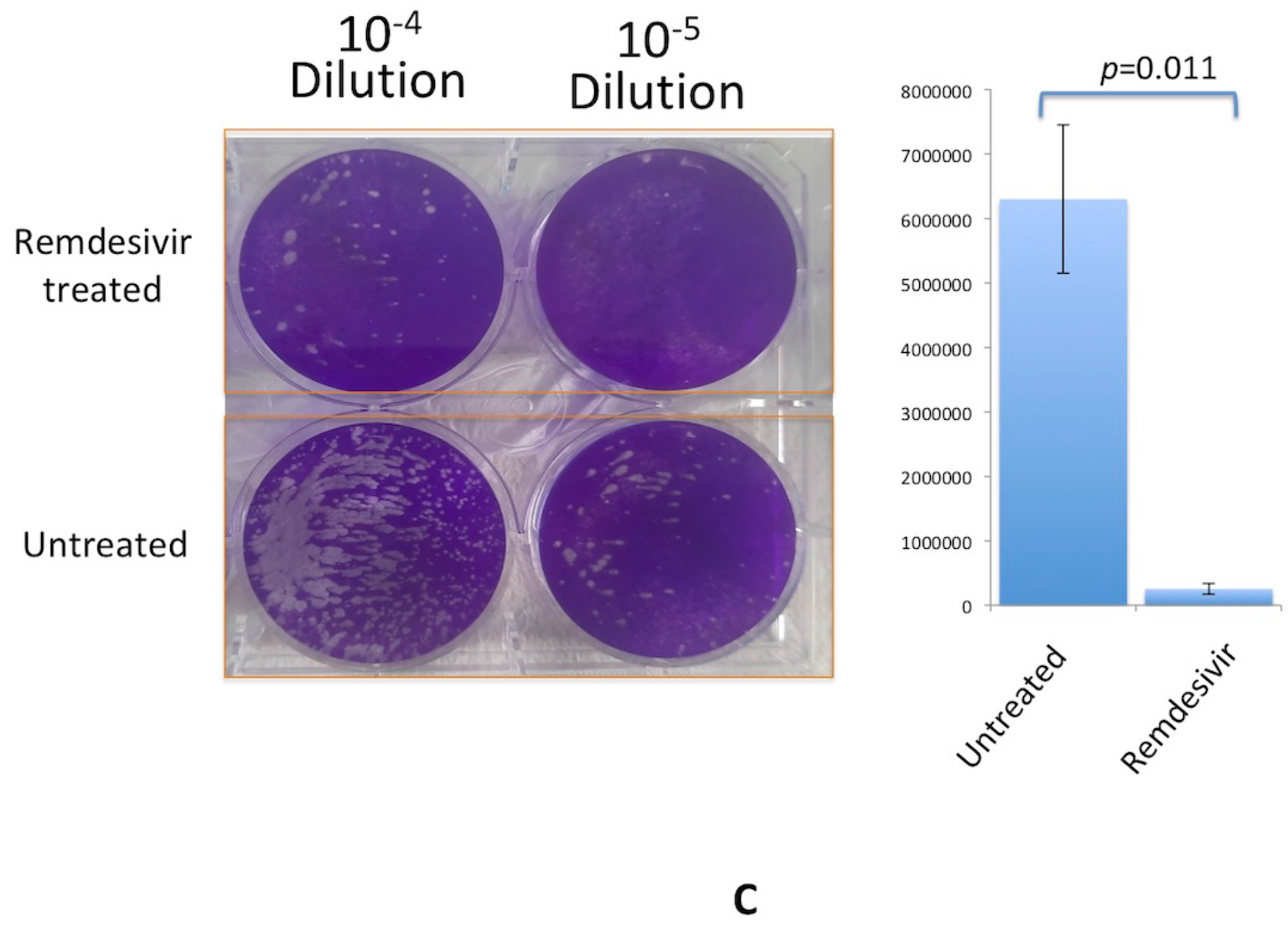
Effects of remdesivir on CHPV replication. **1A**. Cytopathic effects in cultured vero E6 cells infected with CHPV in presence of (bottom) and in absence of remdesivir (middle). Uninfected cells treated with remdesivir were also shown (top). **1B**. Western blot analysis of CHPV infected, remdesivir treated and untreated cells and corresponding supernatants. Western blot was done with rabbit anti CHPV-N antibody. As an internal control beta-tubulin antibody was used. **1C**. Plaque assay to determine quantitative difference in the CHPV production in the supernatants of remdesivir treated and untreated cells. 10^-4^ and 10^-5^ dilutions of supernatants were used.

We have also performed plaque assay to confirm above results and also to determine the number of virus particles produced in the CHPV infected vero cell supernatants in presence and absence of remdesivir. We observed significant reduction (∼95%, *p* value= 0.011) in the number of plaques when cells are treated with 1mM remdesivir compared to that in untreated cells. Average number of virus produced in 1mM remdesivir treated cell supernatants were 2.5 × 10^5^, whereas that number was 6.3×10^6^ in drug untreated cells (Fig. 1C).

### CHPV replication in inhibited by AZT and nevirapine

In addition to remdesivir, we have also examined the effects of HIV-1 reverse transcriptase inhibitor drugs AZT and nevirapine on CHPV replication. Infection of CHPV in vero cells treated with AZT and nevirapine showed that both the drugs inhibit virus replication but inhibitory effects are lesser than remdesivir. We did not observe major difference in cytopathic effects in 1mM drug tretaed and CHPV infected cells as compared to drug untreated and infected cells (Data not shown). However, analysis of cell lysates from 1 mM drug treated and CHPV infected cells showed lower levels of CHPV N proteins compared to untreated cells (Fig. 2A and 2B, Top). This results indicated that 1 mM concentrations of both AZT and Nevirapine had inhibitory effects on CHPV replication. AZT and nevirapine also inhibited virus production at 1 mM concentration of the drugs as shown by western blot of viral proteins obtained from cell free supernatants (Fig. 2A and 2B; middle panel). These results were further confirmed by plaque assay. Averge number of plaques produced are 3.3×10^6^ (∼50% inhibition, *p* value = 0.017) and 1.9×10^6^ (∼66% inhibition, *p* value =0.026) in AZT and nevirapine treated cells respectively as compared to ∼6.3× 10^6^ plaques in untreated cells (Fig. 2C).

**Figure 2.**
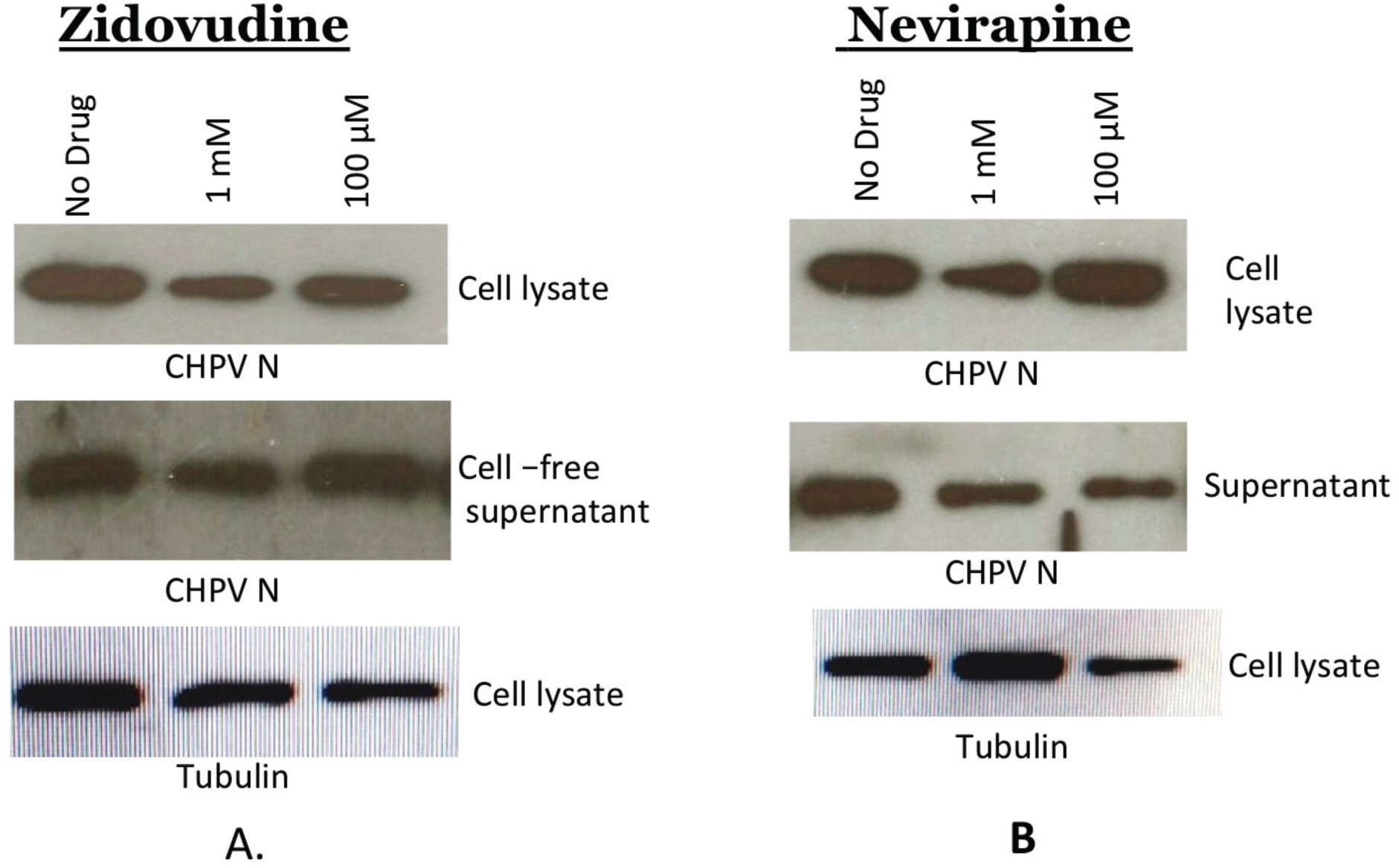
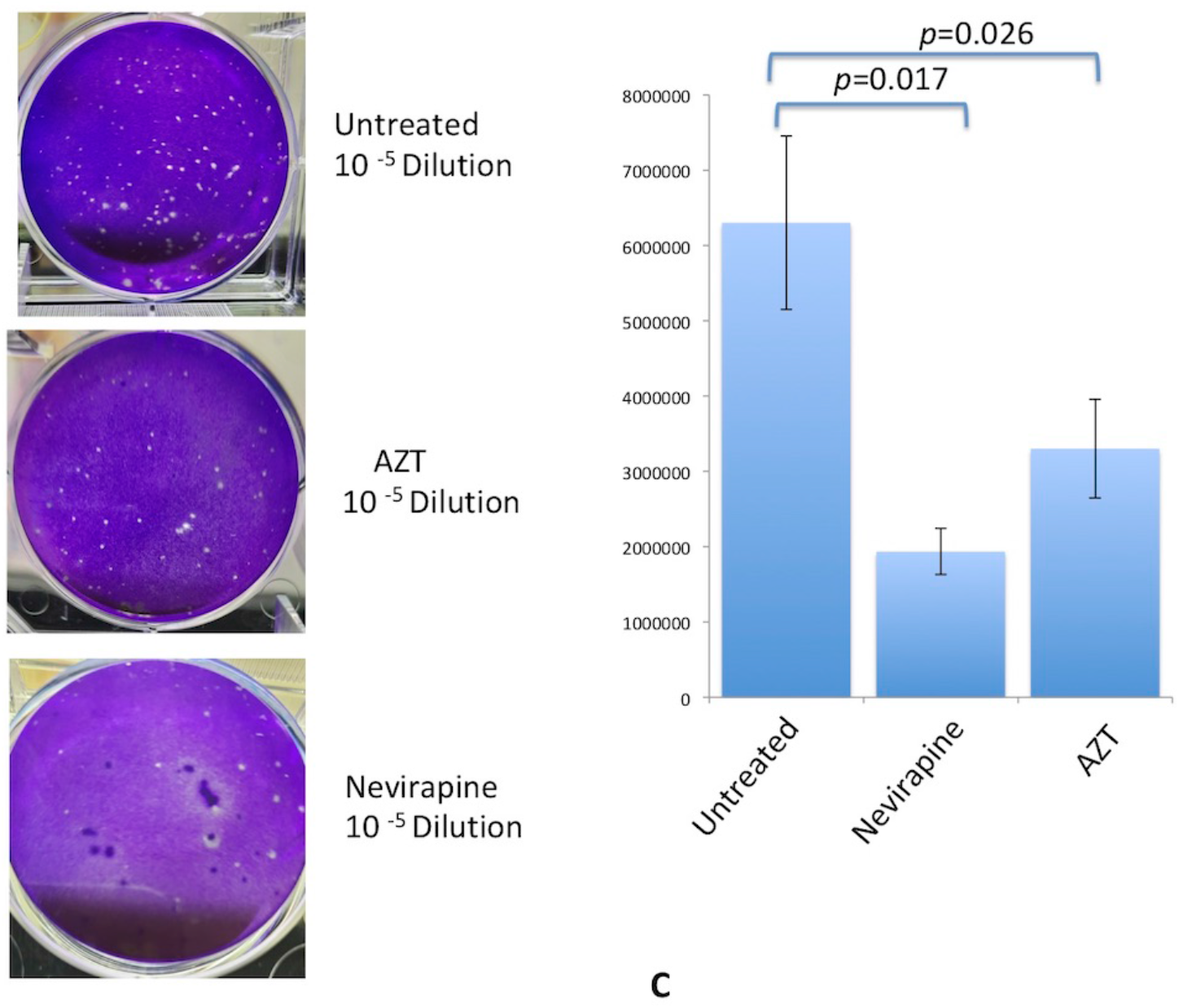
Effects of AZT and nevirapine on CHPV replication. **2A**. Western blot analysis of CHPV infected, AZT treated and untreated cells and corresponding supernatant. Concentrations of AZT used were indicated. Western blot was done with rabbit anti CHPV-N antibody. As an internal control beta-tubulin antibody was used. **2B**. Western blot analysis of CHPV infected, nevirapine treated and untreated cells and corresponding supernatants. Concentrations of nevirapine used were indicated. Western blot was done with rabbit anti-CHPV-N antibody. As an internal control beta-tubulin antibody was used. **2C**. Plaque assay to determine quantitative differences in the virus production in the supernatants of AZT and nevirapine treated and untreated cells. Numbers of plaques obtained at 10^---5^ dilutions of respective supernatants were graphically presented.

### Remdesivir, AZT and nevirapine inhibit the synthesis of CHPV transcripts

To see whether remdesivir inhibits CHPV transcripts synthesis, we extrated total RNA from remdesivir treated cells and prepared cDNA. From cDNA, we amplified G sequence. End point PCR products after 25 cycles of amplification were analyzed on agarose gel. Visible differences were observed in 1.6 kb amplicon intensity in the remdesivir treated lane compared to undreated lanes. (Fig. 3A). As an internal control we used GAPDH primers to amplify the ∼200 bp product and no visible differences was noticed in remdesivir treated and untreated lanes (Fig. 3A). We did not observe any difference in band intensity of amplified G gene in case of AZT and nevirapine (Data not shown).

**Figure 3.**
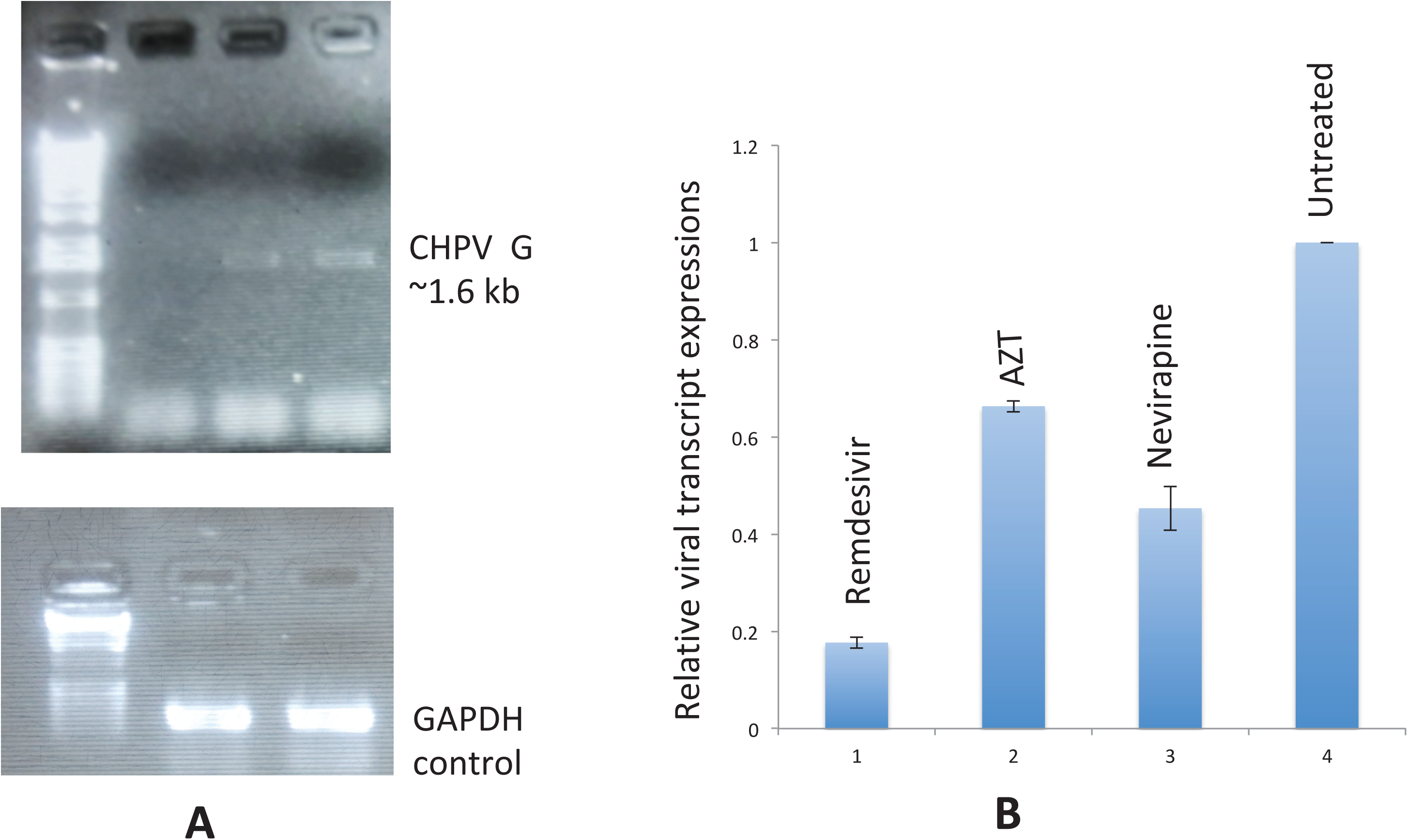
Effects of remdesivir, AZT and nevirapine on CHPV transcription. **3A**. Agarose gel electrophoresis of PCR amplified G gene of CHPV (top) and GAPDH gene (bottom) from remdesivir treated and untreated vero E6 cells. **3B**. Real-time PCR analysis of CHPV RNA transcripts synthesized in remdesivir, AZT and nevirapine treated and CHPV infected vero E6 cells. Levels of transcripts synthesized were compared with untreated cells. We compared the levels of G gene products in drug treated vs. untreated cells normalized with GAPDH levels. Relative levels of CHPV G gene transcripts were calculated and compared using 2^−ΔΔCt^ formula.

We next performed real-time PCR assay to measure the quantitative decrease of RNA transcripts synthesis in the presence of remdesivir. We also wanted to know by real-tinme PCR if AZT and nevirapine treatment of vero cells affected CHPV transcription even at a smaller extent as western blotting showed lesser N protein cell lysate and supernatants. Methodology of real-time PCR was described in materials and methods section. We measured the CHPV envelope gene transcripts synthesized using CHPV G specific primers and as internal control we have measured GAPDH. Our results showed that remdesivir treatment resulted about 5.6 fold (∼17%) decrease of treanscript levels compared to untreated cells (Fig. 3B). Nevirapine and AZT treatment caused 2.2 fold and 1.4 fold decrease respectively in the transcript synthesis compared to no drug treatment (Fig. 3B).

### Structure modeling and assessment

BLASTp search against PDB database indicated Vesicular Stomatitis Virus L protein (VSV L) as the most similar sequence (Percentage sequence Identity: 59.79% and E-value: 0). Hence we used the structure of VSV L protein (PDB ID: 6U1X, Length: 2109 AA) as a template to model the 3D structure of CHPV (Length: 2092 AA). Alignment showed motif F of finger subdomain of both the viruses has close to 99 % similarity which predicts identical rNTP binding efficiency. Polymerase active domains in the palm sequence are also identical and both the polymerases have GDNQ sequence in the active sites, which is also a signature of rabhdoviruses (Fig. 4A). Distribution of Φ and ψ torsion angles in Ramachandran Plot showed presence of 89.4% of residues in the core region, 10.2% in allowed regions and only 0.4% of residues in disallowed regions (Table 2). Analysis of non-bonded interactions between different atom types using ERRAT server showed an average score of 92.75 for modeled structure that indicated highly accurate structure models. Together, distribution of Φ and ψ angles in Ramachandran plot and high ERRAT score indicated the quality of predicted 3-D structures as being reliable and within the acceptable range of different stereo-chemical features (Table 2). Superimposition of modeled structure of CHPV L with the VSV L structure showed a root mean square deviation (RMSD) of 0.171 Å, which indicated absence of any significant structure deviation between the modeled and templated structures (Fig. 4B). This further confirmed the reliability of the modeled structure.

**Figure 4.**
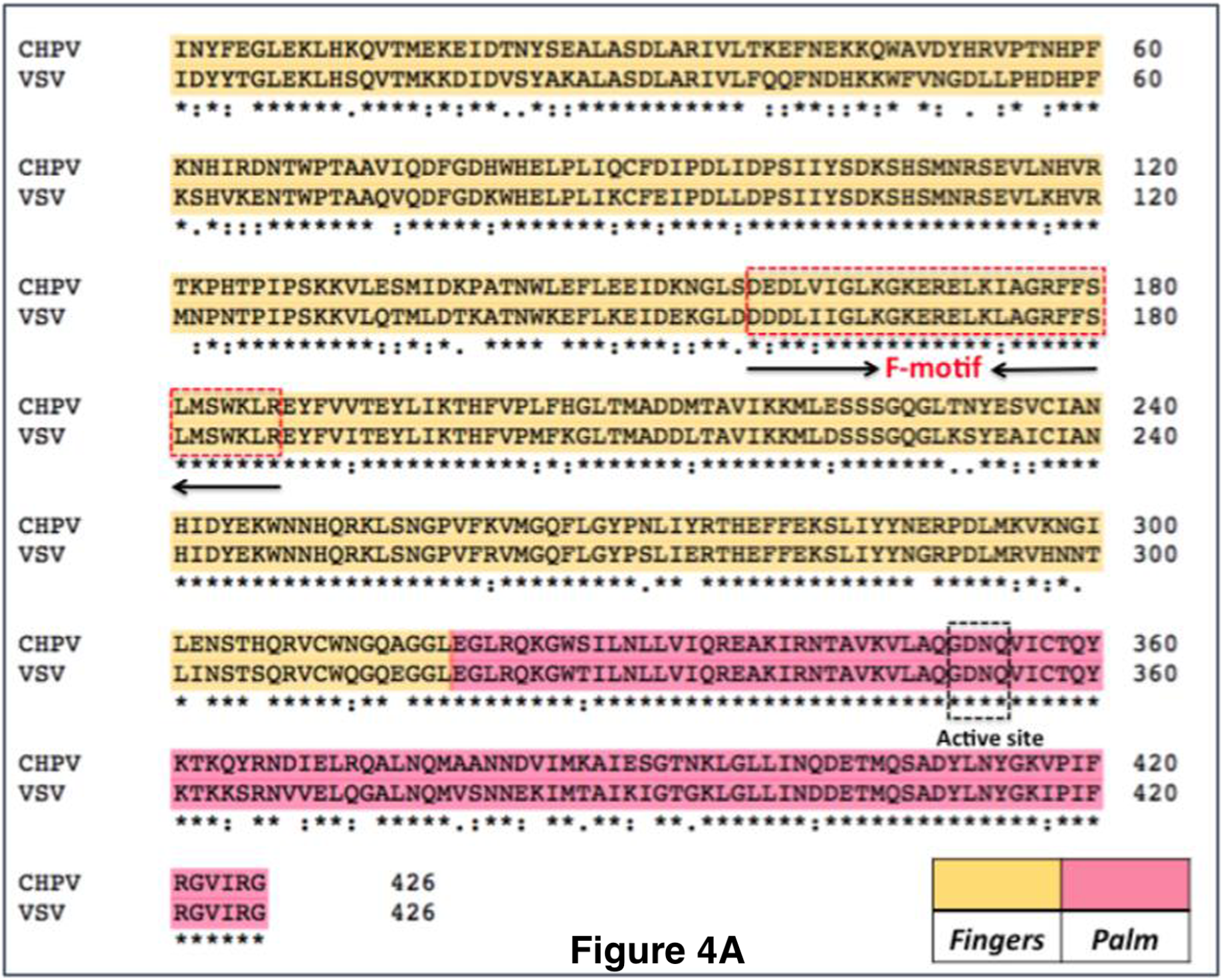
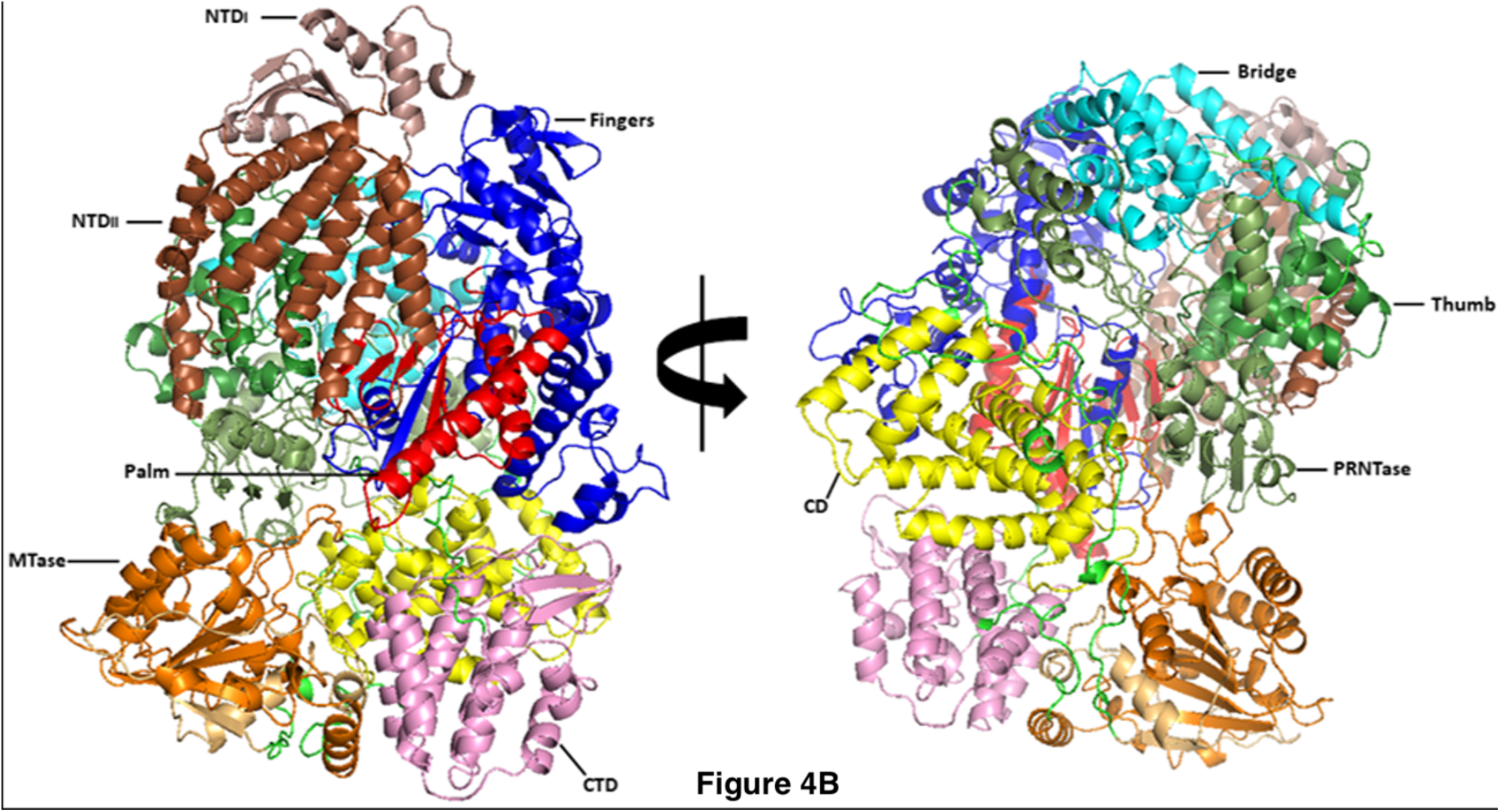
Predicated 3D structure of CHPV polymerase protein L. **4A**. Sequence alignment of the RdRp domain (fingers and palm regions) of CHPV and VSV L proteins. **4B**. Predicted 3D structure of CHPV L protein. Two views of predicted 3D structure of CHPV L protein showed in cartoon form with different colors and labeled according to domain and subdomains (left) and the 180° rotation view (mirror image) of the CHPV L protein (right) depicted to get a clear view of invisible domains and subdomains. N-terminal (NTDs), Methyl transferase (MTase), PRNTase and Thumb domains are marked.

### Docking of antiviral drugs with CHPV L proteins

#### A. Interaction with nevirapine

The following eight bonds were observed between nevirapine and CHPV L protein out of which one was hydrogen bond (K66) and remaining seven were hydrophobic contacts (W68, S204, D464, F206, D217, G205, R218) (Table 3). The CHPV L protein– nevirapine interaction had a binding energy and X-score of −4.21 kcal mol−1 and −7.18 kcal mol−1 respectively which indicated strong binding affinity.

**Table 3:** Docking studies of CHPV with remdesivir, nevirapine and AZT.

| **Drug** | **Binding Energy (kcal/mol)** | **X-score (kcal/mol)** | **Number of Interactions** | **Hydrogen Bonds** | **Hydrophobic contacts** |
| --- | --- | --- | --- | --- | --- |
| Remdesivir | -4.56 | -7.72 | 12 | P555, V697 | L556, H558, E2086, K691, Q687, T695, A690, A696, K698, F2084 |
| Nevirapine | -4.21 | -7.18 | 8 | K66 | W68, S204, D464, F206, D217, G205, R218 |
| AZT | -3.81 | -6.90 | 10 | S2087, H2089, R1136, G579 | S587, N584, L582, I2085, S578, S1138 |

The nevirapine interacting residues of L, identified by docking analysis, are in the N-terminal domian of the CHPV polymerase (Fig. 5 and Fig. 6). Recent studies indicated that this region is important in transcription and this also interact with the residues in th active site of the polymerase in the folded structure.

**Figure 5:**
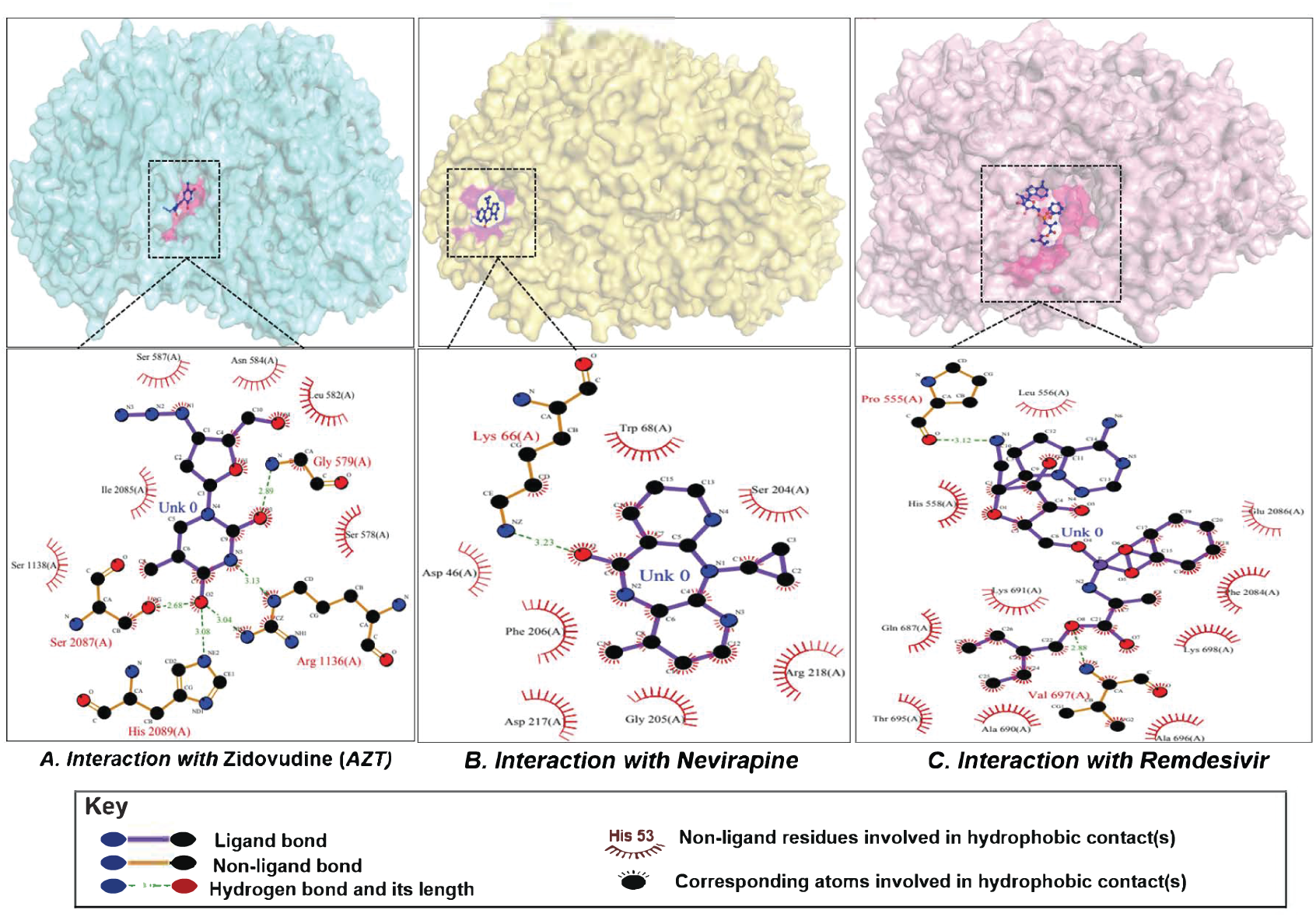
Spatial position of each of three drug on CHPV protein and details of interacting residues between each drug and CHPV protein. Drug-interacting amino acids of CHPV protein is highlighted in pink and drugs are highlighted in blue. Detailed information of interacting residues is given in table 3.

**Figure 6.**
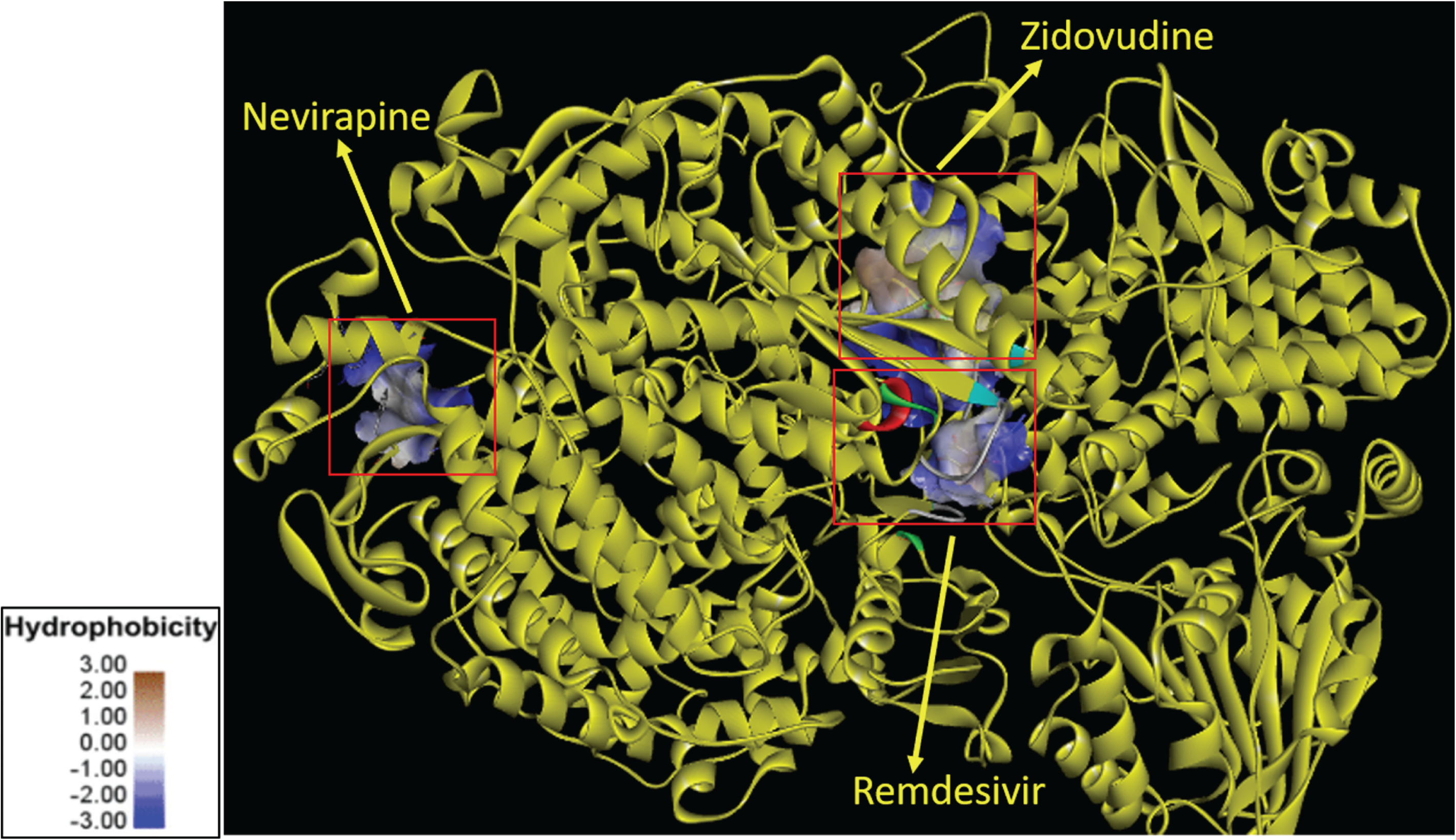
Surface hydrophobicity of CHPV L protein with remdesivir, zidovudine and nevirapine. The figure highlights the comparative binding sites and hydrophobicity of the interacting residues of the above-mentioned drugs to the CHPV protein. The color of the surfaces represents the level of hydrophobicity. The blue, white, and brown colors represent low, mediate, and high hydrophobicity, respectively. Protein structures are drawn using Discovery studio 4.0.

#### B. Interaction with remdesivir

Remdesivir is a well known broad-spectrum antiviral drug which is a nucleotide prodrug of an adenosine analog. It binds to the viral RNA-dependent RNA polymerase and inhibits viral replication by terminating RNA transcription prematurely. It was observed that remdesivir interacts with CHPV L protein by forming the highest number of bonds (Table 3). Analysis of remdesivir-CHPV L protein complex showed P555, V697 were involved in hydrogen bonding and L556, H558, E2086, K691, Q687, T695, A690, A696, K698, and F2084 of CHPV L protein were bound by hydrophobic interactions. The docking score was −4.56 kcal mol−1 and its corresponding X-score was −7.72 kcal mol−1. The X-score was found to be lowest for remdesivir among all three drugs, which confirms that remdesivir interacts with CHPV L protein with high affinity. This high binding affinity comes from its 12 interactions that are present between remdesivir and CHPV L protein (Table 3). Residues Phe 555, L556 and H 558 are located in the finger subdomain of the the polymerase and K691,Q687,T695,A690, A696, K898 are in the palm domain, whereas E2086 and F2084 are in C-terminal domain. Residues in the finger domains are important in selecting and binding to the incoming NTPs. Palm region has the polymerase active site and residues in this region are important stabilizing the template and newly synthesized chain and its translocation (Fig. 5).

#### C. Interaction with AZT

AZT was observed to interact with CHPV L by forming a total of ten bonds (**Table 3**). Out of a total 10 bonds four were hydrogen bonds that included G579, R1136, S2087 and H2089 and six hydrophobic contacts namely, S587, N584, L582, I2085, S578 and S1138 (Fig 5). The CHPV L protein–AZT interaction had the binding energy of −3.81 kcal mol−1 and X-Score of −6.90 kcal mol−1. The results showed that AZT formed the maximum number of hydrogen bonds as compared to other two drugs.

These overall results reveal that remdesivir has high binding affinity compared to other drugs which has an even lower binding affinity and polymerase binding sites of these drugs are also different (Fig. 5 and Fig. 6). Further experimental MIC and TEM data may strengthen the conclusions drawn based on bioinformatics analysis.

## Discussions

In the present study we have investigated the effects of remdesivir, AZT and nevirapine on CHPV replication with an interest to see if any of these drugs can be of clinical use in treating CHPV associated disease and to prevent death. These three are previously known polymerase inhibitor drugs with successful clinical use. Remdesivir is now in use against EBOV and SARS-CoV-2 (Emergency use), whereas AZT and nevirapine are included in HAART regime in treating HIV/AIDS. We were intrigued to find out that remdesivir significantly inhibited CHPV replication in vero E6 cells. In silico model obtained from molecular docking study suggested strong binding of remdesivir with CHPV polymerase enzyme L (binding energy -4.56 kcal/mol and x-score -7.72 kcal/mol)(Table 3). Both the vero cell based *ex vivo* study and docking based modeling thus supported our hypothesis about remdesivir’s role in binding to CHPV RdRp and inhibiting the virus replication. A closer view of docked remdesivir with CHPV L showed that remdesivir binds to finger and palm subdomain of the polymerase (Fig. 5 and 6). As a nucleoside analog, remdesivir is predicted to cause chain terminaton and inhibit the entry of ATP into active site of the RdRp where polymerase adds the incoming nucleotide to the 3’ termini of growing RNA chain. Molecular docking revealed that remdesivir interacts with six residues in the palm region (K691, Q687,T695, A690, A696 and K698) and these in addition to residues in the finger region makes a hydrophobic pocket for this inhibitor with favorable free energy. Recent *in vitro* study with purified EBOV and RSV RdRp showed that remdesivir completes with ATP and causes chain termination at +5 position ahead of primer termini (i+6 position) **(30)**. *In vitro* studies with purified CHPV polymerase will further help in better understanding the mechanism of remdesivir mediated inhibition of CHPV replication. AZT, another drug used in this study, binds in a region close to remdesivir binding site in the finger domain but with lesser affinity and less favorable free energy (Table 3). Inhibitory effects of AZT was expected to be lesser than remdesivir as AZT is a nucleotide analog and we observed the expected trend in vero cells infecetd with CHPV when treated with this drug. Nevertheless, these results are important in further in silico designing of new NTP analog drugs similar to AZT and targeted to the same binding pocket.

To our knowledge this study is the first to examine the inhibitory effects of nevirapine on a non-segmented negative stranded RNA virus and based on our results this drug may be studied against other negative stransded viruses in future. We have found that the drug inhibits CHPV and binds to the N-terminal domain of the polymerase L. A recent study have shown that D236 and E290 residues in N-terminal dmain of VSV L is important in efficient viral RNA trnscription **(31)**. These two residues are conserved in CHPV L and nevirapine interacts with the amino acids which are in close proximity of these residues. These findings may help in designing new classes of small molecules which will target this region of the polymerases belonging to this virus family. Moreover, non-nucleoside inhibitors are of better choice as antivirals because of lesser toxicity as compared to NRTIs as evident from clinical studies on treatment of HIV/AIDS **(32)**. In summary, we were able to show that remdsivir may be a targeted drug for CHPV. Our results thus hold promise to conduct further studies in animal model and clinical use of remdesivir against the CHPV disease. A drug cocktail consisting of redesivir, nevirapine and AZT may be a better option to prevent CHPV infection and associated death.

## Aknowledgement

We thank Prof. Sanjeev Sinha, Department of Internal Medicine, AIIMS, New Delhi for providing AZT and Nevirapine used in this study. We are thankful to Prof. Alo Nag, Department of Biochemistry, University of Delhi South Campus (UDSC) and Dr. Jagreet Kaur, Department of Genetics, UDSC for sharing reagents and instrument facility.

## References

1. Jacobo-Molina A, Ding J, Nanni RG, Clark AD Jr, Lu X, Tantillo C, Williams RL, Kamer G, Ferris AL, Clark P, et al. Crystal structure of human immunodeficiency virus type 1 reverse transcriptase complexed with double-stranded DNA at 3.0 A resolution shows bent DNA. Proc Natl Acad Sci U S A. 1993 Jul 1;90(13):6320–4. doi: 10.1073/pnas.90.13.6320. PMID: 7687065; PMCID: PMC46920.

2. Kirchdoerfer, R.N., Ward, A.B. Structure of the SARS-CoV nsp12 polymerase bound to nsp7 and nsp8 co-factors. Nat Commun 10, 2342 (2019). https://doi.org/10.1038/s41467-019-10280-3

3. Liang B, Li Z, Jenni S, Rahmeh AA, Morin BM, Grant T, Grigorieff N, Harrison SC, Whelan SPJ. Structure of the L Protein of Vesicular Stomatitis Virus from Electron Cryomicroscopy. Cell. 2015 Jul 16;162(2):314–327. doi: 10.1016/j.cell.2015.06.018. Epub 2015 Jul 2. PMID: 26144317; PMCID: PMC4557768.

4. Picarazzi F, Vicenti I, Saladini F, Zazzi M, Mori M. Targeting the RdRp of Emerging RNA Viruses: The Structure-Based Drug Design Challenge. Molecules. 2020 Dec 3;25(23):5695. doi: 10.3390/molecules25235695. PMID: 33287144; PMCID: PMC7730706.

5. Jin Z, Wang Y, Yu XF, Tan QQ, Liang SS, Li T, Zhang H, Shaw PC, Wang J, Hu C. Structure-based virtual screening of influenza virus RNA polymerase inhibitors from natural compounds: Molecular dynamics simulation and MM-GBSA calculation. Comput Biol Chem. 2020 Apr;85:107241. doi: 10.1016/j.compbiolchem.2020.107241. Epub 2020 Feb 26. PMID: 32120300.

6. van Hemert FJ, Zaaijer HL, Berkhout B. In silico prediction of ebolavirus RNA polymerase inhibition by specific combinations of approved nucleotide analogues. J Clin Virol. 2015 Dec;73:89–94. doi: 10.1016/j.jcv.2015.10.020. Epub 2015 Nov 5. PMID: 26587786.

7. Gharbi-Ayachi A, Santhanakrishnan S, Wong YH, Chan KWK, Tan ST, Bates RW, Vasudevan SG, El Sahili A, Lescar J. Non-nucleoside Inhibitors of Zika Virus RNA-Dependent RNA Polymerase. J Virol. 2020 Oct 14;94(21):e00794–20. doi: 10.1128/JVI.00794-20. PMID: 32796069; PMCID: PMC7565630.

8. Basak S, Mondal A, Polley S, Mukhopadhyay S, Chattopadhyay D. Reviewing Chandipura: a vesiculovirus in human epidemics. Biosci Rep. 2007 Oct;27(4-5):275–98. doi: 10.1007/s10540-007-9054-z. PMID: 17610154; PMCID: PMC7087735.

9. Menghani S, Chikhale R, Raval A, Wadibhasme P, Khedekar P. Chandipura Virus: an emerging tropical pathogen. Acta Trop. 2012 Oct;124(1):1–14. doi: 10.1016/j.actatropica.2012.06.001. Epub 2012 Jun 18. PMID: 22721825.

10. Mishra AC. Chandipura encephalitis: a newly recognized disease of public health importance. In: Mishra AC, editor. National Institute of Virology Commemorative Compendium. Golden Jubilee Publications, Pune, India: NIV; 2004. p. 1–20

11. Bhatt PN, Rodrigues FM. Chandipura: a new Arbovirus isolated in India from patients with febrile illness. Indian J Med Res. 1967 Dec;55(12):1295-305. PMID: 4970067.

12. Rao BL, Basu A, Wairagkar NS, Gore MM, Arankalle VA, Thakare JP, et al. A large outbreak of acute encephalitis with high fatality rate in children in Andhra Pradesh, India, in 2003, associated with Chandipura virus. Lancet 2004; 364 : 869–74.

13. Gurav YK, Tandale BV, Jadi RS, Gunjikar RS, Tikute SS, et al. (2010) Chandipura virus encephalitis outbreak among children in Nagpur division, Maharashtra, 2007. Indian J Med Res 132:395–399.

14. Tandale, BV., Tikute, SS., Arankalle, VA., Sathe, PS. and Joshi, MV. 2008. Chandipura virus: a major cause of acute encephalitis in children in North Telangana, Andhra Pradesh, India. J. Med. Virol., 80:118–124.

15. Chadha MS, Arankalle VA, Jadi RS, Joshi MV and Thakare JP. 2005. An outbreak of Chandipura virus encephalitis in the eastern districts of Gujarat state, India. Am. J. Trop. Med. Hyg. 73: 566–570.

16. Mavale MS, Fulmali PV, Geevarghese G, Arankalle VA, Ghodke YS, Kanojia PC, Mishra AC. Venereal transmission of Chandipura virus by Phlebotomus papatasi (Scopoli). Am J Trop Med Hyg. 2006 Dec;75(6):1151-2. PMID: 17172384.

17. Te Velthuis AJW, Grimes JM, Fodor E. Structural insights into RNA polymerases of negative-sense RNA viruses. Nat Rev Microbiol. 2021 May;19(5):303–318. doi: 10.1038/s41579-020-00501-8. Epub 2021 Jan 25. Erratum in: Nat Rev Microbiol. 2021 Feb 2;: PMID: 33495561; PMCID: PMC7832423.

18. Warren TK, Jordan R, Lo MK, Ray AS, Mackman RL, Soloveva V, Siegel D, Perron M, Bannister R, Hui HC, Larson N, Strickley R, Wells J, Stuthman KS, Van Tongeren SA, Garza NL, Donnelly G, Shurtleff AC, Retterer CJ, Gharaibeh D, Zamani R, Kenny T, Eaton BP, Grimes E, Welch LS, Gomba L, Wilhelmsen CL, Nichols DK, Nuss JE, Nagle ER, Kugelman JR, Palacios G, Doerffler E, Neville S, Carra E, Clarke MO, Zhang L, Lew W, Ross B, Wang Q, Chun K, Wolfe L, Babusis D, Park Y, Stray KM, Trancheva I, Feng JY, Barauskas O, Xu Y, Wong P, Braun MR, Flint M, McMullan LK, Chen SS, Fearns R, Swaminathan S, Mayers DL, Spiropoulou CF, Lee WA, Nichol ST, Cihlar T, Bavari S. Therapeutic efficacy of the small molecule GS-5734 against Ebola virus in rhesus monkeys. Nature. 2016 Mar 17;531(7594):381-5. doi: 10.1038/nature17180. Epub 2016 Mar 2. Erratum in: ACS Chem Biol. 2016 May 20;11(5):1463. PMID: 26934220; PMCID: PMC5551389.

19. Higgs ES, Gayedyu-Dennis D, Fischer Ii WA, Nason M, Reilly C, Beavogui AH, Aboulhab J, Nordwall J, Lobbo P, Wachekwa I, Cao H, Cihlar T, Hensley L, Lane HC. PREVAIL IV: A Randomized, Double-Blind, 2-Phase, Phase 2 Trial of Remdesivir vs Placebo for Reduction of Ebola Virus RNA in the Semen of Male Survivors. Clin Infect Dis. 2021 Nov 16;73(10):1849–1856. doi: 10.1093/cid/ciab215. PMID: 33709142.

20. Hoenen T, Groseth A, Feldmann H. Therapeutic strategies to target the Ebola virus life cycle. Nat Rev Microbiol. 2019 Oct;17(10):593–606. doi: 10.1038/s41579-019-0233-2. Epub 2019 Jul 24. PMID: 31341272.

21. Santoro MG, Carafoli E. Remdesivir: From Ebola to COVID-19. Biochem Biophys Res Commun. 2021 Jan 29;538:145–150. doi: 10.1016/j.bbrc.2020.11.043. Epub 2020 Nov 19. PMID: 33388129; PMCID: PMC7836944.

22. Lo MK, Albariño CG, Perry JK, Chang S, Tchesnokov EP, Guerrero L, Chakrabarti A, Shrivastava-Ranjan P, Chatterjee P, McMullan LK, Martin R, Jordan R, Götte M, Montgomery JM, Nichol ST, Flint M, Porter D, Spiropoulou CF. Remdesivir targets a structurally analogous region of the Ebola virus and SARS-CoV-2 polymerases. Proc Natl Acad Sci U S A. 2020 Oct 27;117(43):26946–26954. doi: 10.1073/pnas.2012294117. Epub 2020 Oct 7. PMID: 33028676; PMCID: PMC7604432.

23. Parker WB, White EL, Shaddix SC, Ross LJ, Buckheit RW Jr, Germany JM, Secrist JA 3rd, Vince R, Shannon WM. Mechanism of inhibition of human immunodeficiency virus type 1 reverse transcriptase and human DNA polymerases alpha, beta, and gamma by the 5’-triphosphates of carbovir, 3’-azido-3’-deoxythymidine, 2’,3’-dideoxyguanosine and 3’-deoxythymidine. A novel RNA template for the evaluation of antiretroviral drugs. J Biol Chem. 1991 Jan 25;266(3):1754-62. PMID: 1703154.

24. Mitsuya H, Weinhold KJ, Furman PA, St Clair MH, Lehrman SN, Gallo RC, Bolognesi D, Barry DW, Broder S. 3’-Azido-3’-deoxythymidine (BW A509U): an antiviral agent that inhibits the infectivity and cytopathic effect of human T-lymphotropic virus type III/lymphadenopathy-associated virus in vitro. Proc Natl Acad Sci U S A. 1985 Oct;82(20):7096–100. doi: 10.1073/pnas.82.20.7096. PMID: 2413459; PMCID: PMC391317.

25. Das K, Martinez SE, Bauman JD, Arnold E. HIV-1 reverse transcriptase complex with DNA and nevirapine reveals non-nucleoside inhibition mechanism. Nat Struct Mol Biol. 2012 Jan 22;19(2):253–9. doi: 10.1038/nsmb.2223. PMID: 22266819; PMCID: PMC3359132.

26. Das K, Martinez SE, Bandwar RP, Arnold E. Structures of HIV-1 RT-RNA/DNA ternary complexes with dATP and nevirapine reveal conformational flexibility of RNA/DNA: insights into requirements for RNase H cleavage. Nucleic Acids Res. 2014 Jul;42(12):8125–37. doi: 10.1093/nar/gku487. Epub 2014 May 31. PMID: 24880687; PMCID: PMC4081091.

27. Livak KJ, Schmittgen TD. Analysis of relative gene expression data using real-time quantitative PCR and the 2(-Delta Delta C(T)) Method. Methods. 2001 Dec;25(4):402–8. doi: 10.1006/meth.2001.1262. PMID: 11846609.

28. Colovos, Chris, and Todd O. Yeates. 1993. “Verification of Protein Structures: Patterns of Nonbonded Atomic Interactions.” Protein Science. https://doi.org/10.1002/pro.5560020916.

29. Laskowski, R. A., M. W. MacArthur, D. S. Moss, and J. M. Thornton. 1993. “PROCHECK: A Program to Check the Stereochemical Quality of Protein Structures.” Journal of Applied Crystallography. https://doi.org/10.1107/s0021889892009944.

30. Tchesnokov EP, Feng JY, Porter DP, Götte M. Mechanism of Inhibition of Ebola Virus RNA-Dependent RNA Polymerase by Remdesivir. Viruses. 2019 Apr 4;11(4):326. doi: 10.3390/v11040326. PMID: 30987343; PMCID: PMC6520719.

31. Qiu S, Ogino M, Luo M, Ogino T, Green TJ. Structure and Function of the N-Terminal Domain of the Vesicular Stomatitis Virus RNA Polymerase. J Virol. 2015 Oct 28;90(2):715–24. doi: 10.1128/JVI.02317-15. PMID: 26512087; PMCID: PMC4702691.

32. MacArthur RD, Novak RM, Peng G, Chen L, Xiang Y, Hullsiek KH, Kozal MJ, van den Berg-Wolf M, Henely C, Schmetter B, Dehlinger M; CPCRA 058 Study Team; Terry Beirn Community Programs for Clinical Research on AIDS (CPCRA). A comparison of three highly active antiretroviral treatment strategies consisting of non-nucleoside reverse transcriptase inhibitors, protease inhibitors, or both in the presence of nucleoside reverse transcriptase inhibitors as initial therapy (CPCRA 058 FIRST Study): a long-term randomised trial. Lancet. 2006 Dec 16;368(9553):2125–35. doi: 10.1016/S0140-6736(06)69861-9. PMID: 17174704.

